# The Time Course of Recognition Memory Impairment and Glial Pathology in the hApp-J20 Mouse Model of Alzheimer’s Disease

**DOI:** 10.1101/495960

**Authors:** Kamar E. Ameen-Ali, Julie E. Simpson, Stephen B. Wharton, Paul R. Heath, Paul Sharp, Gaia Brezzo, Jason Berwick

## Abstract

The role of cellular changes in the neurovascular unit is increasingly being investigated to understand the pathogenesis of Alzheimer’s disease. The aim of the current study was to determine the time course of recognition memory impairment in the J20 mouse model of AD, in relation to neuroinflammatory responses and the pathology of Aβ.

Male hAPP-J20 and wild-type mice were assessed at 3, 6, 9, and 12 months of age. The spontaneous object recognition (SOR) task provided a measure of memory, with assessment of both a short delay (1 min) and a long delay (4 hrs). Immunohistochemistry was used to characterise Aβ-deposition, and quantify astrocyte and microglial responses.

At all ages tested J20 mice had impaired long-term, but preserved short-term, recognition memory. Wild-types demonstrated preserved long-term memory up to 9 months of age, and preserved short-term memory at all ages tested. Plaque pathology in the J20 mice was present from 6 months onwards, with co-localisation of reactive microglia and activated astrocytes. Reactive microglia and astrocyte activation in the hippocampus were significantly greater in the J20 mice at 9 months, compared to wild-types.

This study contributes to our understanding of the pathological and cognitive mechanisms at play in AD. J20 mice showed impairment in retaining information over longer periods from an early age, preceding the deposition of Aβ and glial activation. Defining early physiological changes in relation to cognitive decline could provide insight into new therapeutic targets early in the disease process, when intervention is most likely to effectively slow disease progression.

## Introduction

Alzheimer’s disease (AD) is characterised by a decline in cognitive skills, such as memory and visuo-spatial capabilities. Pathological characteristics of AD include beta amyloid (Aβ) plaques, hyperphosphorylated tau forming neurofibrillary tangles [NFTs; 1], synapse and neuron loss, and neuroinflammation [2]. However, the mechanistic links between cognitive phenotypes and specific neuropathologies needs further definition.

A number of AD mouse models have been generated to recapitulate the pathological and behavioural features of the disease (see [3, 4, 5]). No one model yet exists that fully recapitulates all aspects of AD; however, each model enables specific aspects of the disease to be interrogated.

The hAPP-J20 mouse model of AD includes both the Swedish (K670N and M671L) and Indiana (V7171F) mutations on a C57Bl/6 x DBA2J background. This model is particularly useful for investigating amyloid pathology, as it features an overexpression of Aβ1-42, with early plaque deposition from around five months of age. Previous cognitive tests have revealed varying results, but generally report impairments in recognition and spatial memory that closely coincide with amyloid deposition in the hippocampus and cortex, though this is dependent on task parameters [3, 6–10].

The J20 mouse model develops pathological hallmarks typical of AD, such as astrocyte activation and reactive microglia. Astrocytes are part of the neurovascular unit and play a role in maintaining homeostasis [11]. When activated, astrocytes undergo morphological changes, such as developing shorter and thicker processes, and releasing pro-inflammatory factors [12]. Microglia are also reactive in AD [13, 14]; both astrocytes and microglia respond to Aβ plaque formation, but more research is needed to determine whether they have a neuroprotective function, or contribute to the progression of neurodegeneration [15]. Microglial responses are complex and diverse, and may be differentially associated with AD pathology, with a decrease in microglial motility reducing neuroprotection, but an increase in phagocytosis contributing to the progression of neurodegeneration [16]. In the J20 mouse model, increases in microglial reactivity and activated astrocytes relative to wild-type controls have been reported as a function of age [17], although reports have also found astrocytes with morphological changes with a decrease in number and cell complexity [6].

Recognition memory is the ability to recognise when something has been previously encountered, and is a form of declarative memory that also includes episodic memory (memory for a past experience in your life; [see 18–23]) which is impaired in AD. The spontaneous object recognition (SOR) task [24] has been successfully used to assess recognition memory in rodents following selective brain impairment, with further task adaptations allowing for more complex forms of memory involving spatial and contextual discriminations [25–28]. Lesion studies have revealed that for the standard SOR task, the perirhinal cortex is necessary for accurate recognition [29–33], but not the hippocampus or the fornix (subcortical fibre pathway leading to the hippocampus) [28, 30, 34]. When the delay between the sample and test phase is substantially long, studies have, however, reported impairments on the SOR task in rodents with hippocampal lesions [35, 36]. The increased delay diminishes the memory strength for the familiar object, making successful discrimination during the test phase more difficult [37].

Previous research has shown that J20 AD mice are able to perform successfully on the SOR task up to 15–16 months of age, with a delay between sample and test phases of up to one hour [38]. If the delay is increased to four hours, the J20 mice become impaired at a younger age of 2–3 months [9] or six to eight months [7] depending on the exact task protocol. If the delay is further increased to 24 hours, the J20 mice are unable to discriminate between novel and familiar objects from as young as four months of age [8]. Although it appears that J20 mice experience impairments in recognition memory both as they age, and with an increased demand on memory through longer delays, it is unclear how much of this impairment can be attributed to age, the length of the delay, and the disease progression.

The aim of the current study was to determine the time course of recognition memory impairment in the J20 mouse model of AD, in relation to neuroinflammatory responses and the pathology of Aβ, to elucidate whether these pathologies precede cognitive impairment or occur later. Such information is likely to be informative as to what should be modulated in order to affect cognition in AD.

Two SOR tasks were performed; the first involved a short delay between sample and test phases of one minute (short-term recognition memory), and the second involved a long delay of four hours (long-term recognition memory). Comparing performance on these two tasks enabled a characterisation of the decline of separate memory processes in normal aging (via performance of the wild-type controls), and in disease (via the J20 AD mice) at distinct time points in order to define the temporal relationship of pathological cellular changes to these cognitive defects.

## Method

### Subjects

Male hemizygous transgenic mice (hAPP-J20) that overexpress a mutant form of the human amyloid precursor protein (hAPP) carrying the Swedish (K670N and M671L) and Indiana (V7171F) mutations were obtained from Jackson Laboratories (Bar Harbor, ME), and crossed with C57Bl/6J female mice to produce hAPP-J20 mice and wild-type littermate controls. Pups were weaned at 21 days of age and progeny ear-clipped and genotyped (Transgene Forward 5’-GGT GAG TTT GTA AGT GAT GCC-3’, Transgene Reverse 5’-TCT TCT TCT TCC ACC TCA GC-3’).

An a priori power analysis yielded a suitable total sample size of 45 for obtaining statistically meaningful results. A total of 54 male hAPP-J20 and wild-type mice were used, with eight transgenic mice for each of the four time points (with the exception of the 3 month and 9 month groups which had seven), and six wild-type mice for each of the four time points (3 months, 6 months, 9 months, and 12 months of age). The higher allocation of animals in the transgenic groups was due to potential attrition from spontaneous death occurring before three months of age, likely due to seizures as the J20 model is known for abnormal neural hyperexcitability [39].

All animals were housed in groups of between two and five (based on keeping littermates together, regardless of genotype) in diurnal conditions (12-h light-dark cycle) with testing carried out during the light phase. Water was available ad libitum throughout the study. All animals were food deprived to 90% of the free-feeding body weight of age matched controls throughout testing. Mean weights were as follows: 3 month wild-type = 27g; 3 month J20 = 24g; 6 month wild-type = 29g; 6 month J20 = 27g; 9 month wild-type = 37g; 9 month J20 = 30g; 12 month wild-type = 39g; 12 month J20 = 31g. All experiments were performed in accordance with the U.K. Animals (Scientific Procedures) Act (1986) and associated guidelines, and have been reported in accordance with the ARRIVE (Animal Research: Reporting of *In Vivo* Experiments) guidelines.

### Apparatus

Object recognition memory was tested in a square shaped arena which included an outer corridor surrounding the perimeter (Figure 1). The apparatus was 50cm long and 50cm wide, with the inner open field being 34cm long and 34cm wide. Four mechanical doors (8cm wide) divided the inner open field with the outer corridor, with one door in the centre of each of the four sides of the arena. These doors could be opened and closed remotely, and separate removable corner walls divided each of the four corridors surrounding the inner open field. Only one door (and subsequently one corridor) was used in this study for the animals to shuttle between the areas during and between trials, as the corridor served purely as a holding area for the animal in between phases when objects were being changed. Objects were placed centrally in the open field, with one object on the left and one on the right, approximately 2cm or more away from the walls to allow the animals to fully explore the objects.

**Fig. 1.**
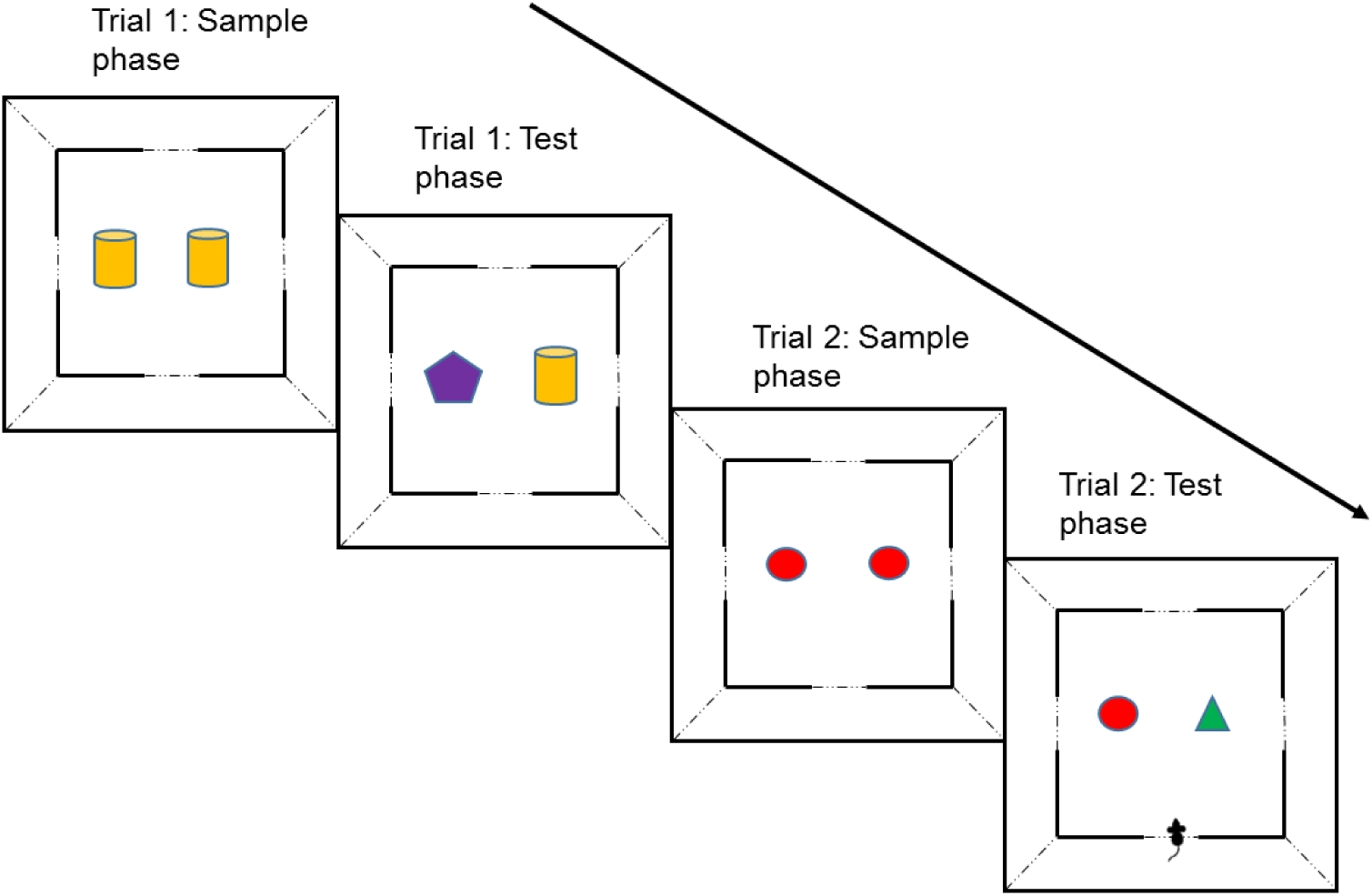
An illustration of the short-term spontaneous object recognition (SOR) memory test procedure showing sample and test phases for two trials with a one minute delay between phases. Each mouse was given a single testing session of 10 trials from which mean D2 scores were derived. For the long-term SOR task with a four hour delay between sample and test phases, a standard one-trial a day version of the task was used with the sample phase occurring in the morning, and the test phase in the afternoon (always at the same time). Multiple trials were carried out on consecutive days and mean D2 scores derived from these raw scores.

### Objects

Various objects of different sizes, shape, colour, and texture were used as stimuli in the object recognition experiments, with identical duplicate objects used within each trial. Each animal did not re-encounter the same object within an object recognition experiment or on any subsequent object recognition test.

### Pretraining

All animals were given two sessions of handling and two sessions of habituation to the testing room, where they remained in a holding cage with their home cage mates for a period of 10 minutes per session. Each group of animals housed together had their own holding cage so as to minimise stress from mixing smells of animals from different home cages.

Pretraining lasted for approximately five days, and involved the completion of five phases aimed to habituate the animals to the environment and task procedure. Phase 1 involved placing the animals in the apparatus to freely explore with their cage mates for 30 minutes. For Phase 2, the animals were individually placed into the apparatus for 20 minutes to freely explore. This was repeated for Phase 3 for 10 minutes. Phase 4 was aimed at training the animals to shuttle between the outer corridor and the open arena. Three 10 minute sessions were carried out and involved placing single MLab rodent pellets (45mg, TestDiet; Richmond, Indiana, USA) in both the centre of the open field, and behind the door leading to the outer corridor. The food was replenished after the completion of each shuttle. Phase 5 introduced objects into the open field, whereby individually the animals shuttled into the open field from the outer corridor and were exposed for three minutes to two identical objects, each baited with a single food pellet. The door leading to the outer corridor was then opened for the animals to shuttle through to the outer corridor, which contained a single food pellet. Once the objects had been changed, the door opened for the animals to shuttle back into the open field containing a duplicate copy of the object from the sample phase and a novel object, again each baited with a single food pellet. After three minutes, the animals then shuttled back in to the outer corridor when the door was opened. This procedure was carried out for a total of three different pairs of objects (not re-used in the experiment proper).

### Test protocol for short-term object recognition memory

Each mouse was given a single testing session of 10 trials (Figure 1). At the start of each session, the animal was placed in the outer corridor (only one of the four were used, and this was the same for all mice), with the door opening immediately so they could move through to the open field. Each animal spent three minutes in the open field exploring the objects during the sample phase. The door to the outer corridor was then opened, and the animal shuttled through to the outer corridor to obtain a single food pellet. After one minute the door opened to allow the animal back into the open field for the test phase which contained a duplicate copy of the now familiar object from the sample phase and a novel object (trial 1). Following three minutes, the door opened and the animal shuttled through to the outer corridor. After a period of one minute, the door was then opened for trial 2 allowing the animal back into the open field for the sample phase of trial 2, and so on. This procedure continued for a total of 10 trials. All objects were baited with a food pellet to encourage exploration of objects without differentially rewarding choices.

The location of the novel object was counterbalanced to minimise bias for left or right exploration within each testing session and also between animals. Objects were also counterbalanced between animals for which was novel and which was familiar, to minimise bias for a particularly salient object. A trial was ended if the animal failed to shuttle to the next area of the apparatus after a period of three minutes, ceasing the testing session.

### Test protocol for long-term object recognition memory

The protocol for long-term object recognition memory followed the week after testing for the short-term object recognition memory had been completed. The test protocol for this task was identical to that of the short-term object recognition memory task, however the delay between sample and test phases on each trial was extended to four hours, in which the animal was removed from the apparatus and placed back in their home cage. Subsequently, it was only possible to carry out one trial a day for each animal, of which four trials were carried out in total on consecutive days.

### Behavioural analysis

Object exploration was defined as when the nose of the animal was <1cm from the object, or if the object was touched with the animal’s nose (directed within 45° of the object) or paws. Sitting or climbing on the object were actions not counted as exploration. Duration of exploration was measured off-line using a computerised stop-watch mechanism whilst exploration was observed by the experimenter on a recording. Discrimination was measured using D2 scores [19] which are calculated using the difference in exploration time (exploration time for the novel object minus the exploration time for the familiar object) divided by the total exploration time. A D2 score was calculated for each individual trial then averaged to give a mean D2 score for each animal, which were then used in the data analysis. The D2 index ranged from −1 to +1, with −1 representing total exploration of the familiar object, +1 representing total exploration of the novel object, and 0 being indicative of no object preference.

### Immunohistochemistry

The animals were humanely euthanased with i.p. injections of sodium pentobarbital (100mg/kg, Euthatal, Merial Animal Health Ltd). They were then transcardially perfused with 0.9% phosphate buffered saline (warmed to 37°C) with the addition of heparin (0.1ml/500ml). Brains were removed from the skull, dissected to separate the two hemispheres, with the right hemisphere subdissected coronally into four parts (cerebellum, frontal lobe, and the remaining tissue cut in half) and then snap frozen in liquid nitrogen for storage at −80°C. The left hemisphere post fixed in 4% paraformaldehyde (0.1M, pH 7.4) and subdissected into four regions, as described, after 24 hours. Following fixation, the tissue was embedded in paraffin wax.

Serial sections (5μm) were cut from the paraffin-embedded tissue. Immunohistochemistry was carried out using a standard avidin-biotin complex (ABC) method. Sections were first deparaffinised, rehydrated to water, followed by quenching of endogenous peroxidase activity through placing the sections in 0.3% H_2_O_2_/methanol for a total of 20 minutes at room temperature (RT).

Sections underwent antigen retrieval based on prior antibody optimisation; single and dual immunolabelling for Iba1 (Ionized calcium binding adaptor molecule 1) underwent microwave antigen retrieval for 10 minutes (pH9), and single and dual immunolabelling for GFAP (glial fibrillary acidic protein) underwent pressure cooker antigen retrieval for 20psi at 120°C for 45s (pH6.5). All immunolabelling for Aβ included additional pre-treatment of the sections in 70% formic acid for 10min at RT.

Following antigen retrieval, sections were incubated in 1.5% blocking solution for 30 minutes at RT, before further incubation in the primary antibody (anti-Iba1 – 1:200, Abcam, UK; biotinylated anti-Aβ – 1:500, BioLegend, USA; anti-GFAP – 1:500, Dako, USA) made in blocking solution for 60 minutes at RT. Antibody binding was visualised using the standard horseradish peroxidase avidin-biotin complex method (Vectastain Elite kit, Vector Laboratories, UK), with 3,3-diaminobenzidinetetrahydrochloride (DAB) as the chromogen (Vector Laboratories, UK; brown). Sections were further counterstained with haematoxylin (Cell Path, UK), dehydrated, and cleared in xylene before mounting in DPX (Leica, UK). Negative controls, with the omission of the primary antibody and isotype controls, were included with every run.

For dual-staining experiments, GFAP or Iba1 expression was visualised as described above. Sections were then incubated with the avidin-biotin blocking kit (Vector Laboratories, UK), according to the manufacturer’s instructions. The tissue was incubated overnight at 4°C with anti-Aβ and visualised with the alkaline-phosphatase-conjugated avidin-biotin complex and alkaline phosphatase substrate 1 (Vector Laboratories, UK, red). Sections were counterstained with haematoxylin.

### Regions of interest

Quantification of GFAP and Iba1 immunoreactivity was carried out using a Nikon Eclipse Ni-U microscope and Nikon DS-Ri1 camera with NIS-Elements BR 4.20.01 64-bit microscope imaging software (Nikon, UK), capturing bright-field images in subregions of the hippocampus and cortex, using a 20x objective. Regions of interest were identified with reference to the mouse brain anatomy atlas by Paxinos and Franklin [40]. For the perirhinal and somatosensory cortices, two adjacent belt transects from the outer cortex through to the white matter border were taken for each animal at 20x magnification. Each belt transect contained up to four images, and the area coordinates for the captured images were taken between AP −1.07 to −2.45 relative to bregma for the somatosensory region, and between AP - 1.67 to -2.53 relative to bregma for the perirhinal cortex. Subfields of the hippocampus (CA1, CA3, and dentate gyrus) were imaged between AP −1.67 to −3.51 relative to bregma. All slides were imaged blind in a random order.

### Neuropathological quantification and statistical analysis

The images were thresholded and the GFAP or Iba1 immunoreactive area of the field determined per total area examined, using the Analysis ^D software. The average percentage area of immunopositive staining was used for statistical analysis for both antibodies, for wild-type and J20 mice, and statistically analysed using a two-way ANOVA with *age* (3, 6, 9, and 12 months) and *genotype* (J20 and wild-type) as between subject factors. All images were analysed blind in a randomised order.

## Results

### Behavioural results: J20 mice show short-term recognition memory up to 12 months of age

Five animals died before reaching the age for behavioural testing; one from the three month J20 group, one from the six month J20 group, one from the nine month J20 group, and two from the 12 month J20 group.

To determine whether the animals performed above chance, a one-sample t-test (two-tailed) was used to compare the mean D2 scores against zero for each genotype group at each of the four ages. Both J20 and wild-type mice could successfully discriminate between novel and familiar objects at all ages tested, though J20 mice did not perform quite as well as wild-type mice (see Figure 2a legend). For the mean D2 scores, there was a significant main effect of genotype (F(1,41) = 6.736, p = 0.013), with the J20 mice overall obtaining lower mean D2 scores compared to wild-types (t(47) = 2.639, p = 0.011), but no significant main effect of age (F(3,41) = 1.728, p = 0.176), or interaction (F(3,41) = 0.329, p = 0.804) was found.

**Fig. 2.**
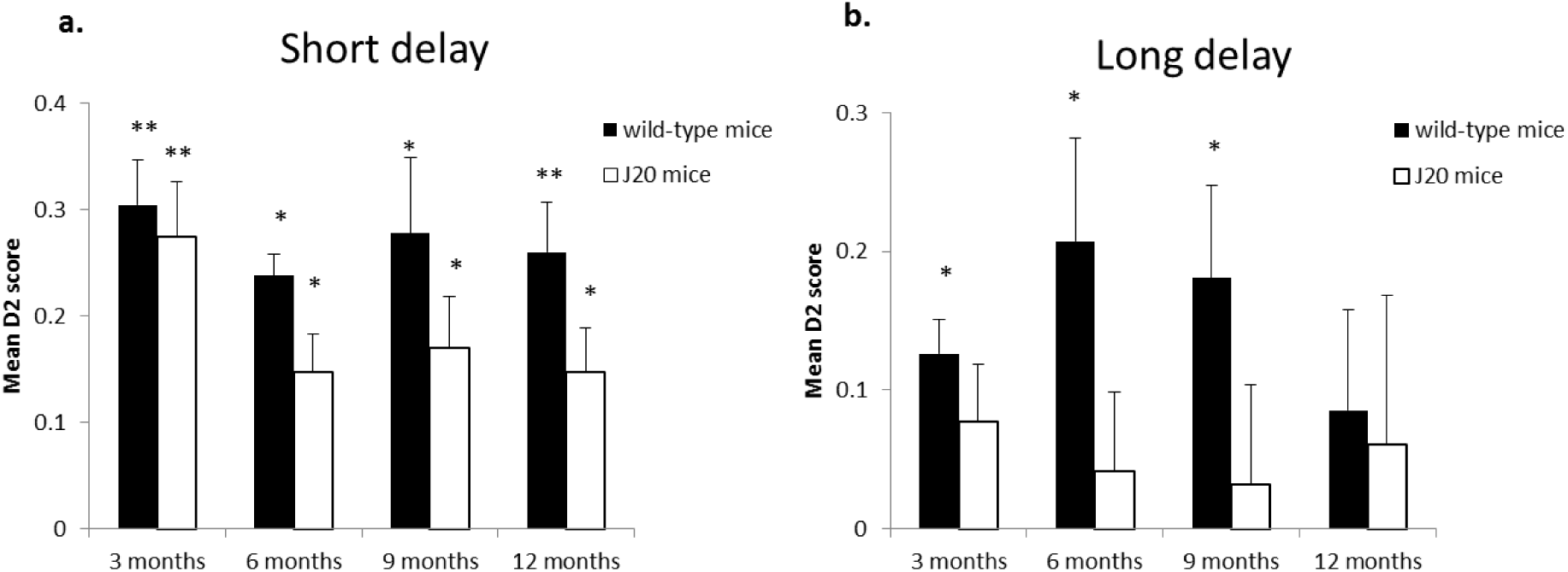
SOR memory tasks. Mean D2 scores from the SOR task with a short delay (a). All the groups significantly explored the novel stimuli more than the familiar stimuli (WT3m: t(5) = 7.225, p = 0.001; WT6m: t(5) = 12.134, p = <0.001; WT9m: t(5) = 3.873, p = 0.012; WT12m: t(5) = 5.501, p = 0.003; Tg3m: t(5) = 5.319, p = 0.003; Tg6m: t(6) = 4.205, p = 0.006; Tg9m: t(5) = 3.554, p = 0.016; Tg12m: t(5) = 3.554, p = 0.016); Mean D2 scores from the SOR task with a long delay (b). The wild-type mice were able to successfully discriminate between the novel and familiar stimuli up to nine months of age, but failed recognition at 12 months of age (WT3m: t(5) = 3.986, p = 0.010; WT6m: t(5) = 2.797, p = 0.038; WT9m: t(5) = 2.720, p = 0.042; WT12m: t(5) = 1.168, p = 0.295). The J20 mice were unable to discriminate at all the ages tested (Tg3m: t(5) = 1.855, p = 0.0123; Tg6m: t(6) = 0.740, p = 0.487; Tg9m: t(5) = 0.452, p = 0.670; Tg12m: t(5) = 0.570, p = 0.594). Independent samples t-tests between J20 and wild-type mice at each age for both tasks were all non-significant (all p = >0.05). Vertical bars show the standard error of the mean. (*) p < 0.05; (**) p < 0.01.

For the total amount of time spent exploring the objects in the test phase, there was a significant main effect of genotype (F(1,41) = 6.329, P = 0.016), with the J20 mice spending significantly less time exploring overall in the test phase compared to wild-types (t(33.265) = 2.263, p = 0.30), and age (F(3,41) = 3.151, p = 0.035), with a general decline in exploration with age. There was no significant interaction between genotype and age (F(3,41) = 2.392, p = 0.082).

### J20 mice are significantly impaired at long-term recognition memory

The wild-type mice were able to successfully discriminate between the novel and familiar stimuli up to nine months of age, but failed recognition at 12 months of age, and the J20 mice were unable to discriminate at all ages tested (see Figure 2b legend). There was a trend towards a significant main effect of genotype on the mean D2 score (F(1, 41) = 3.996, p = 0.052) with the J20 mice overall obtaining lower mean D2 scores compared to wild-types (t(47) = 2.098, p = 0.041), but no main effect of age (F(3, 41) = 0.197, p = 0.898) or significant interaction (F(3, 41) = 0.537, p = 0.659) was found.

For the total amount of time spent exploring the objects in the test phase, there was a significant main effect of genotype (F(1,41) = 20.781, p = <0.001), with the J20 mice spending significantly less time exploring overall in the test phase compared to wild-types (t(38.783) = 4.348, p = <0.001), but no significant main effect of age (F(3,41) = 2.371, p = 0.084), or interaction (F(3,41) = 1.034, p = 0.387) was found.

### Neuropathological results: Astrogliosisis associated with age and not genotype in the hippocampus

Two animals died after behavioural testing before they could be perfused; one from the three month J20 month, and one from the six month J20 group. In addition, the quality of tissue for some regions of interest was not suitable for image analysis, so the group numbers may vary throughout the immunohistochemistry data analysis.

GFAP specifically immunolabelled astrocyte cell bodies and proximal processes in all cases. Although there was no significant difference overall between J20 and wild-type mice, there was a significant difference between these groups at 9 months of age in the hippocampus as a whole (Figure 3a), with astrocytes in J20 mice exhibited a hypertrophied appearance, indicative of astrogliosis (Figure 4a). Negative controls did not show any immunoreactivity.

**Fig. 3.**
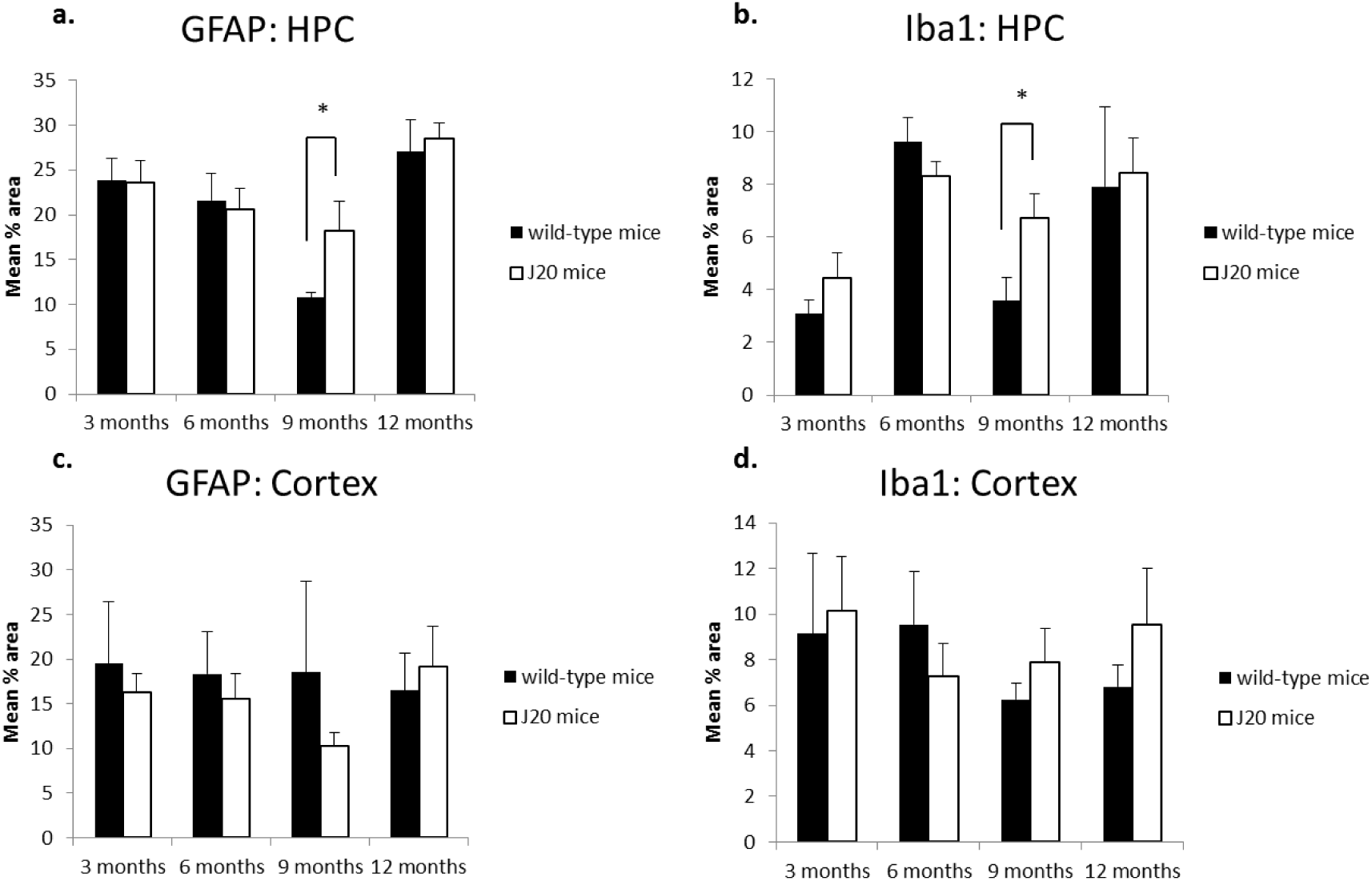
Pooled mean % immunolabelled area. GFAP in the hippocampus (a). J20 mice had significantly greater GFAP expression in the hippocampus at 9 months of age (t(6) = 2.822, p = 0.030), but at no other time point (all p = >0.700); Iba1 in the hippocampus (b). J20 mice had significantly greater Iba1 expression in the hippocampus at 9 months of age (t(6) = 3.169, p = 0.019), but at no other time point (all p = >0.256); GFAP in the cortex with no significant association with either age or genotype (c); Iba1 in the cortex with no significant association with either age or genotype (d). Vertical bars show the standard error of the mean. (*) p < 0.01.

**Fig. 4.**
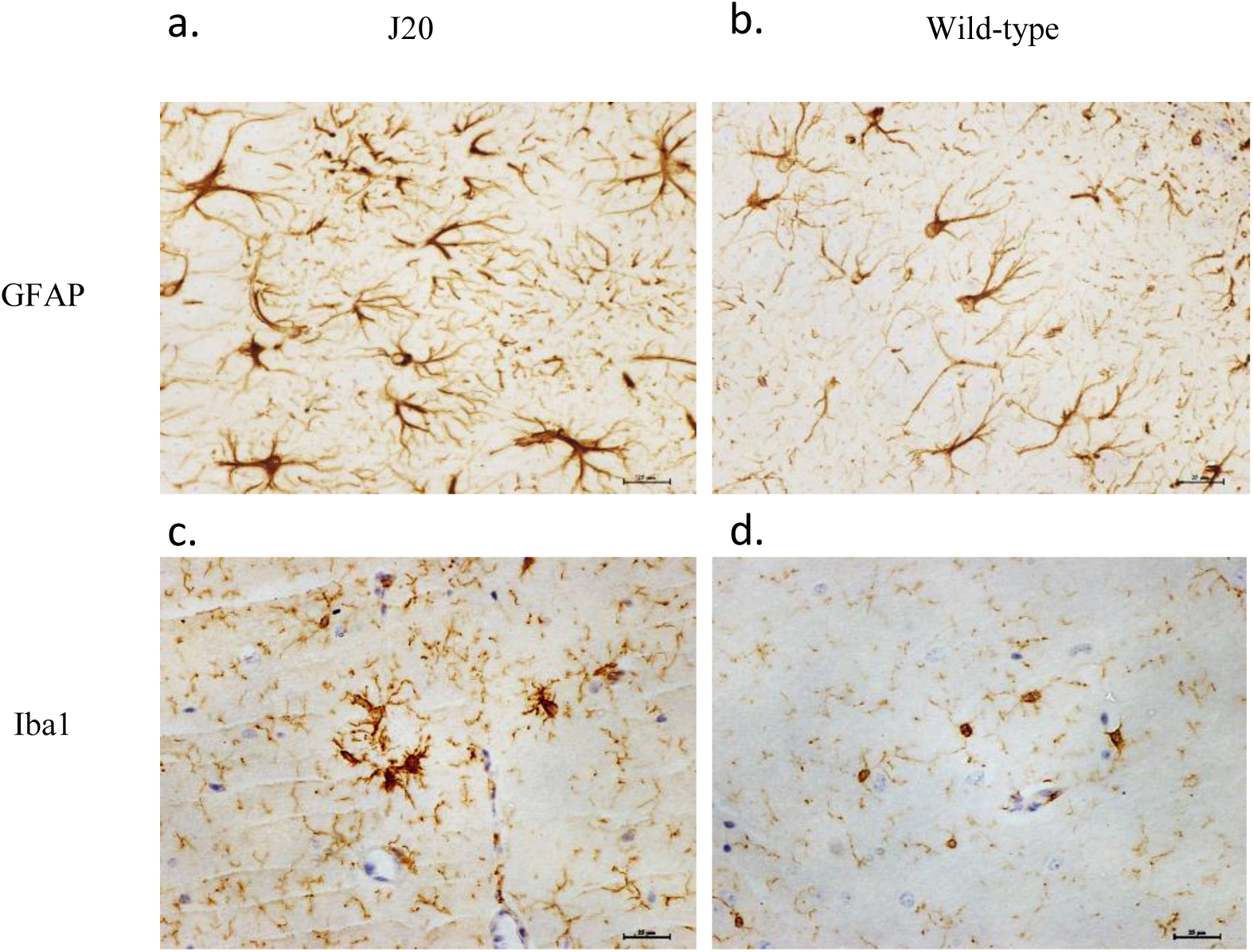
GFAP^+^ astrocytes and Iba1+ microglia in the hippocampal subregion CA1 of 9 month old hAPP-J20 and wild-type mice. At 9 months of age, significant differences between J20 and wild-type mice were found, with both astrocytes and microglia showing greater immunoreactivity in the hippocampus with notable morphological changes. (a) Astrocytes with a prominent hypertrophied appearance were a feature of J20 mice, compared to (b) wild-type mice. Similarly, at 9 months of age (c) microglia in J20 mice displayed a more reactive profile with enlarged cell bodies, compared to (d) wild-type mice. Scale bar represents 25μm. Images are contrast enhanced.

GFAP expression was quantified in the subfields of the hippocampus (CA1, CA3, and dentate gyrus [DG]). GFAP expression significantly varied with age in the CA1 (F(3,26) = 12.117, p = <0.001), CA3 (F(3,28) = 3.091, p = 0.043) and the DG subregions (F(3,30) = 5.633, p = 0.003). Pairwise comparisons revealed significant differences in the immunoreactive profile of GFAP between ages 3 and 6 months (CA1: p = 0.035), 3 and 9 months (CA1: p = <0.001), 6 and 12 months (CA1: p = 0.018), and 9 and 12 months (CA1: p = <0.001; CA3: p = 0.035; DG: p = 0.002). There was no significant association with genotype, and no significant interaction between age and genotype in any subregion (all p = >0.179).

Taking the hippocampal region as a whole, combining the data from the subregions, GFAP expression was significantly associated with age (F(3,31) = 8.102, p = <0.001), but there was no significant association with genotype, and no significant interaction between age and genotype (both p = >0.311). Pairwise comparisons revealed significant differences in the immunoreactive profile of GFAP between 3 and 9 months (p = 0.013), and 9 and 12 months (p = <0.001). Irrespective of genotype, levels of GFAP expression in the hippocampus decreased with age until around 9 months, and then increased again when measured at 12 months of age.

Independent-samples t-tests between J20 and wild-type mice at each time point showed J20 mice had significantly greater GFAP expression in the hippocampus at 9 months of age, but at no other time point (see Figure 3a legend).

### Microgliosis associated with age and not genotype in the hippocampus

Iba1 specifically immunolabelled microglial cell bodies and processes in all cases. Although there was no significant difference overall between J20 and wild-type mice, there was a significant difference between these groups at 9 months of age in the hippocampus (Figure 3b), with microglia in J20 mice displayed a hypertrophic profile with larger cell bodies, indicating microglial reactivity (Figure 4c). Negative controls did not show any immunoreactivity.

Iba1 expression was quantified in the subfields of the hippocampus (CA1, CA3, and DG). Iba1 expression significantly varied with age in both the CA1 (F(3,27) = 4.392, p = 0.012) and DG subregion (F(3,30) = 3.414, p = 0.030). Pairwise comparisons revealed significant differences in the immunoreactive profile of Iba1 between ages 3 and 6 months (CA1: p = 0.013), and 6 and 9 months (CA1: p = 0.049), and a trend towards significant difference between ages 3 and 12 months (DG: p = 0.052). No significant association of Iba1 expression with age was found in the CA3 subregion (F(3,28) = 1.073, p = 0.377). There was no significant association with genotype, and no significant interaction between age and genotype in any subregion (all p = > 0.170).

Taking the hippocampal region as a whole, combining data from the subregions, Iba1 expression was significantly associated with age (F(3,31) = 3.584, p =0.025), but no significant association with genotype and no significant interaction between age and genotype (both p = >0.478). Pairwise comparisons revealed a significant difference in the immunoreactive profile of Iba1 between 3 and 6 months (p = 0.045). Irrespective of genotype, levels of Iba1 expression in the hippocampus increased from 3 to 6 months of age, before decreasing at 9 months, and then increasing again at 12 months of age.

Independent-samples t-tests between J20 and wild-type mice at each time point showed J20 mice had significantly greater Iba1 expression in the hippocampus at 9 months of age, but at no other time point (see Figure 3b legend).

### No age or genotype associated changes in GFAP or Iba1expression in the cortex

GFAP and Iba1 expression was quantified throughout the perirhinal and somatosensory cortex but revealed no significant association with either age or genotype, either when the subregions were analysed separately or when taken together as a whole (all p = >0.263).

### Aβ plaque deposition present in J20 mice

Aβ plaques, predominantly located in the hippocampal and cortical regions, were a prominent feature of the J20 mice, but not wild-type control mice (Figure 5). Negative controls did not show any immunoreactivity. At 3 and 6 months of age, diffuse staining of soluble Aβ was localised primarily in the white matter border between the hippocampal and cortical regions. At 9 months of age, compact Aβ plaques were deposited predominantly in the hippocampus, with some beginning to develop in the cortex. By 12 months of age, compact plaques were located across the hippocampus (all subregions) and sparsely across the somatosensory cortex, but little deposition was found in the perirhinal cortex.

**Fig. 5.**
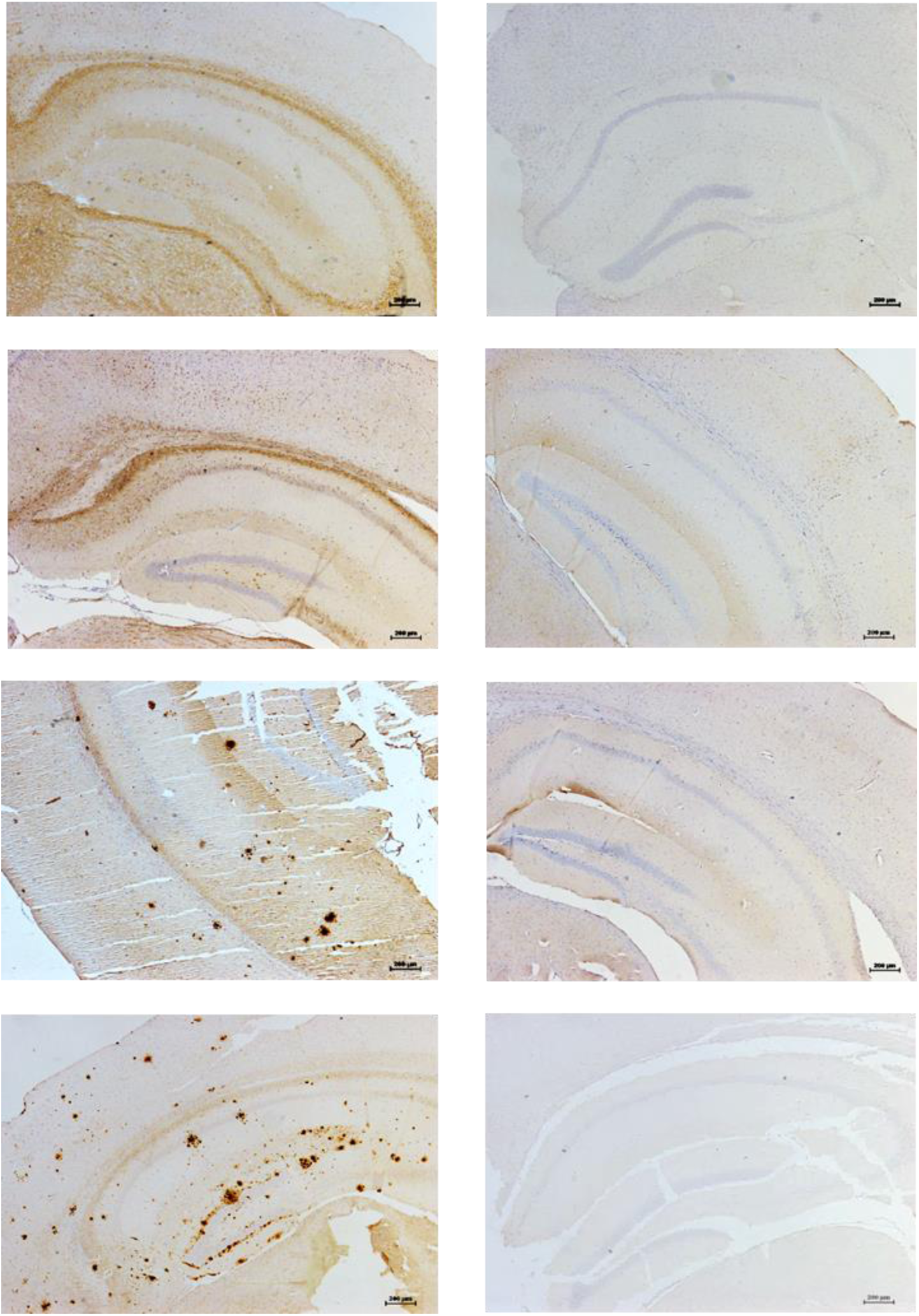
Aβ plaque staining in the hippocampus of hAPP-J20 and wild-type mice. Aβ plaques were predominantly located in the hippocampal and cortical regions in the J20, but not wild-type mice. At 3 and 6 months of age, diffuse staining of soluble Aβ was found in the J20 mice, which progressed to compact Aβ plaques from 9 months of age predominantly in the hippocampus, but also in cortical areas. Scale bar represents 200μm. Images are contrast enhanced.

Dual-labelling of the Aβ plaques and either astrocytes (GFAP) or microglia (Iba1) in the J20 mice displayed co-localisation from 9 months of age (Figures 6a and 6b). Gliotic astrocytes primarily localised with compact plaques, but also infrequently with diffuse plaques. These reactive astrocytes completely surrounded the plaques, with extension of their processes into these deposits. Reactive microglia were intimately associated with both diffuse and compact plaques, with even relatively small depositions surrounded by microglia with large cell bodies and retracted processes.

**Fig. 6.**
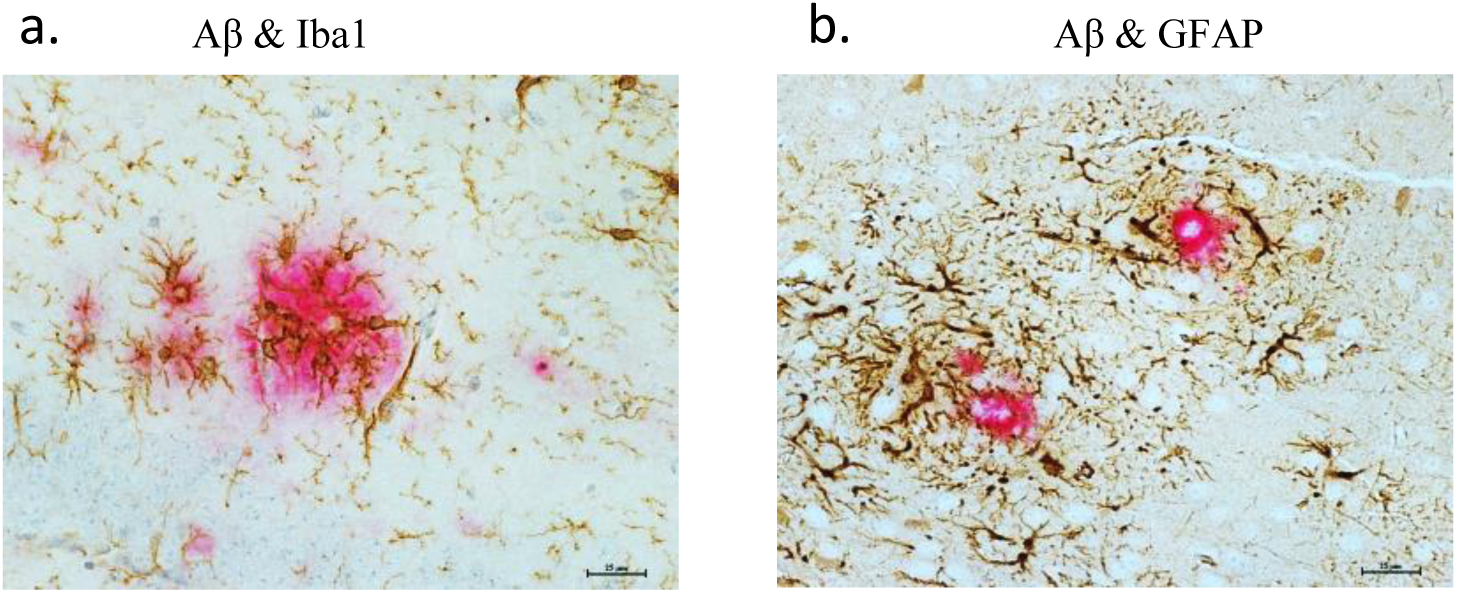
Glial activation in relation to Aβ plaques in the hippocampus of a 12 month old hAPP-J20 mouse. (a) Reactive microglia (brown) with prominent cell bodies were intimately associated with Aβ plaques (red). (b) Reactive astrocytes (brown) were in direct contact with and completely surrounded the plaques (red), with extension of their processes into these deposits. Scale bar represents 25μm. Images are contrast enhanced.

## Discussion

The aim of this study was to determine the time course of recognition memory impairment in the J20 mouse model of AD, in relation to Aβ pathology and neuroinflammatory responses. To our knowledge, this is the first characterisation of Aβ and neuroinflammatory pathology in J20 mice that varies with age, and a dissociation in short- and long-term recognition memory ability that was not observed in wild-type mice.

On a task of short-term recognition memory, both J20 and wild-type mice at all ages tested were able to successfully discriminate between novel and familiar objects. On a task of long-term recognition, however, J20 mice were impaired at all ages tested and wild-type mice were only able to show significant long-term recognition up to 9 months of age. Previous studies have reported that J20 mice are impaired from 4 months of age in a recognition memory task with a 24 hour delay [8], but when the delay is shortened to 4 hours, J20 mice show successful recognition memory up to 6–8 months of age [7]. Harris et al. [9] have reported impairment from 2–3 months. It is clear, therefore, that the length of the delay between the sample and test phase can influence the memory load demand, and the age of the AD mouse (or the progression of the disease), also contributes to the degree to which this demand impacts upon recognition memory performance.

It is important to highlight key methodological differences between the short and long delay recognition memory tasks when comparing performance in the current study. Both tasks were completed within the same apparatus, but the short-term recognition memory task comprised of a 3 minute sample phase and a 3 minute test phase, with trials completed consecutively in a multi-trial paradigm. This paradigm typically results in less variability in the data as animals do not need to be handled repetitively before and after each phase of a trial which can induce anxiety in mice, allowing for multiple trials to therefore be carried out [41]. As it is not possible to perform a long-term recognition memory task with a similar paradigm, the standard version of this task was used with a 10 minute sample phase, and a 5 minute test phase following the 4 hour delay. The longer sample and test phases closely reflected the task used by Harris et al. [9]. Therefore only anecdotal, rather than statistical, comparisons have been made between performances on the two tasks, with a focus on comparing age and genotype differences statistically within each task. The results from these tasks when considered separately, therefore, reflect previous findings from Karl et al. [38] with successful recognition in the short-term SOR task up to 12 months of age, and impairment in long-term SOR from 3 months of age [9]. A recent study, however, has reported impaired recognition memory in J20 mice from six months of age, with a one minute delay between sample and test phases [42]. This contrasts with the results from the current short term SOR task, which J20 mice could perform successfully up to 12 months of age. The difference in results may, in some part, be accountable by task protocol differences whereby the continual trials design in the current study allowed for more trials to be carried out per animal without being handled before and after trial phases. The memory impairment shown by J20 mice in the study by Criscuolo et al. [42] may reflect an inability to perform the task due to confounding factors, such as anxiety, rather than a clear memory deficit.

Immunohistochemistry revealed no quantifiable differences in GFAP and Iba1 expression in the cortical subregions between J20 and wild-type mice, at all ages. Taking the hippocampal region as a whole, GFAP and Iba1 expression varied with age, with GFAP expression decreasing until 9 months of age and then increasing; and Iba1 expression increasing until 6 months of age, decreasing at 9 months, and then increasing again at 12 months. There were no overall significant differences between J20 and wild-type mice; however, GFAP and Iba1 immunoreactivity was significantly higher in J20 mice than in wild-type controls at 9 months of age. Aβ plaques were deposited from 9 months of age in the J20 mice, predominantly in the hippocampus, consistent with previous pathological descriptions of this model [43]. There were no significant neuropathological changes in the perirhinal cortex region, or the cortical region more widely when the results were pooled together. In addition, plaque deposition was relatively sparse in this region. The lack of recognition memory impairment observed in the J20 mice in the short delay task is therefore not surprising, considering this type of memory has been shown to be perirhinal cortex dependant [29–33].

There are clear morphological changes showing astrocyte activation and reactive microglia in the J20 mice (Figure 4a and 4c). There were no significant overall differences between J20 and wild-type mice in terms of GFAP and Iba1 expression in the hippocampus, however there was a significant genotype difference at 9 months of age for both of these glial responses. For GFAP expression, the difference could be attributed to a significant decline at 9 months of age, and for Iba1 expression, the difference could be attributed to a significant increase at 6 months of age, which then reduces again at 9 months. There is a lack of consistency amongst previous studies on glial pathology in aging, with some reporting astrocyte or microglial hyperplasia and hypertrophy [44–47] or, in contrast, atrophy [48]. It is possible that age-related variations in glial response may also be region specific, particularly in terms of morphological changes [49]. In the current study, such age-related changes are not significant in the J20 mice, perhaps due to a more chronic state of astrocyte activation and microglial reactivity that ultimately results in this case in a significant difference in GFAP and Iba1 expression between wild-type and J20 mice at 9 months of age.

The current study did not clearly identify a causal link between the neuropathological and cognitive changes measured, but does provide a characterisation of these changes and when they occur. As previously mentioned, J20 mice up to at least 12 months of age have preserved short-term recognition memory function which is perirhinal cortex dependent [29–33]. Alongside the lack of pathological changes in the cortex across all ages measured, these findings suggest that J20 mice have preserved cortical function. Recent data from our lab supports this assumption, as we found that 9–12 month old J20 mice have preserved haemodynamic and neural responses measured in the somatosensory cortex when compared to wild-type mice [50], contrary to previous studies [51–54]. If J20 mice have preserved cortical function overall, this would explain the lack of short-term memory impairment in J20 mice.

From 3 months of age, J20 mice have impaired recognition memory when there is a long delay between sample and test phases, which studies have previously shown to be hippocampal-dependent [35–36]. This impairment cannot be attributed to significant changes in glial pathology as these were not found at 3 months of age. There was, however, clear diffuse staining of soluble Aβ primarily on the white matter border between the hippocampal and cortical regions that could, alongside a related neuropathological change not measured in this current study, be correlated with the impairment in long-term recognition memory in J20 mice. For instance, oligomeric levels of Aβ were not assessed in the current study, but they do play a key role in the pathogenesis of AD with studies showing that cognitive dysfunction coincides with the presence of significant Aβ oligomers in the Tg2576 AD mouse model [55]. In addition, reducing levels of Aβ oligomers resulted in improved cognition in transgenic mice overexpressing APP [56], and in the 5xFAD and APP/PS1 AD mouse models [57, 58]. It is possible that the long-term recognition memory impairment observed in the J20 mice from 3 months of age could be linked to early oligomer formation. Further work should also characterise synaptic function in the hippocampus which has been reported be impaired in APP mice prior to Aβ plaque deposition [59–62].

Astrogliosis manifests in upregulation of GFAP expression, with hypertrophic cell bodies and processes and astrocyte proliferation [63]. Although astrocyte hypertrophy is a common feature of AD, it may also be important to consider astrocyte atrophy and the associated functional changes, as this has been reported in the early stages of the disease in transgenics, such as the 3xTg-AD model [64–66]. This model exhibits both Aβ and NFTs allowing assessment of astrocyte response to both pathological features, and therefore may recapitulate aspects of AD more effectively than the J20 mouse model which only overexpresses levels of Aβ. A recent study by Goodall et al. [67] reported an overall age-associated increase in astrocyte activation in C57 mice, but in the current study this pattern was not observed and the lack of genotype- or age-associated proliferation may be partially attributable to astrocyte atrophy. Moreover, astrocyte atrophy can lead to synapse loss and reduced neuronal metabolic support, which can ultimately result in early cognitive deficits [66]. As previously mentioned, it would therefore be useful to characterise hippocampal synaptic function particularly to see if this is in part accountable for the long-term recognition memory impairment observed in the J20 mice from 3 months of age.

The current study showed that Aβ compact plaques were co-localised and infiltrated by activated astrocytes and reactive microglia, which has been previously reported [68–70], and Simpson et al. [71] specifically reported a trend to increased GFAP expression with neuritic, rather than diffuse, plaques. Research is ongoing to elucidate whether the activated glia act as a neuroprotective barrier through degrading and clearing Aβ [63, 72], or whether they contribute to the degeneration progression through the release of inflammatory mediators [73]. Research suggests that glial activation is a defensive response against injury, disease, or infection, but excessive activation, or activation sustained over longer periods, may act as a contributing factor to degeneration [74–76]. A recent study by Mathur et al. [77] proposed that astrocytes not only form a protective barrier around amyloid plaques, but astrogliosis negatively related to measures of cognitive impairment, showing a potential neuroprotective role.

It is important to acknowledge that using a single sex in this study may present as a potential limiting factor. Previous research varies in the extent to which sex differences may have an effect on AD phenotypes in mice, particularly in terms of whether certain hormones provide a protective or deleterious effect [3]. Due to the between- and within-subject design of the current study, we opted to exclusively use male mice to minimise variability, but recognise the need for more research into sex differences in disease models and the effect of hormones, particularly in preclinical studies for drug development.

In summary, the current study showed J20 mice were impaired in retaining information over longer periods from an early age, which preceded the deposition of Aβ and glial activation, but were able to retain information over short periods of time, up to at least 12 months of age. The study was limited in terms of the neuropathological markers that were observed, but nevertheless demonstrates that Aβ deposition and cellular neuroinflammatory responses do not precede cognitive impairment. As a clear dissociation in performance on recognition memory was found for the J20 mice, further work to elucidate the pathology of synaptic function would be useful to establish whether this or oligomeric changes may be related to the observed recognition memory deficits and the accumulation of Aβ.

## Acknowledgements

This work was funded by an Alzheimer’s Research UK Interdisciplinary Research Grant (ARUK-IRG2014-10).

